# Genetic mapping in Collaborative Cross mouse strains identifies loci that affect initial sensitivity to cocaine

**DOI:** 10.1101/2025.01.12.629464

**Authors:** SA Schoenrock, CH Gaines, P Kumar, S Khan, J Farrington, MT Ferris, F Pardo-Manuel de Villena, W Valdar, JA Bubier, LM Tarantino

**Affiliations:** University of North Carolina at Chapel Hill; The Jackson Laboratory

**Keywords:** QTL, locomotor, acute, addiction, gene discovery, RI Lines

## Abstract

We identified two Collaborative Cross (CC) strains, CC004/TauUncJ (CC004) and CC041/TauUncJ (CC041), that differ significantly for locomotor response and self-administration of cocaine. In the current study, we crossed each of these strains to C57BL/6NJ (B6N) mice to produce two F2 populations and identify genetic loci that influence locomotor response to cocaine. We identified three significant loci on chromosomes 7, 11 and 14 in the CC041 F2 mapping cross that collectively explain 14% of the phenotypic variance for locomotor response to cocaine. We used a bioinformatic approach to identify high quality candidate genes that are genetically plausible, have functional relevance and are suitable for further exploration. Our study is the first to use CC strains to perform QTL mapping for addiction-related phenotypes and proposes several candidate genes for follow-up analyses.

## INTRODUCTION

Approximately 2.2 million individuals in the United States use cocaine regularly and 1.5 million meet the diagnostic criteria for Cocaine Use Disorder (CUD) (CBHSQ 2016, Abuse 2020). The rate of cocaine use has increased in recent years and cocaine-related overdose deaths have also increased. (John and Wu 2017, Cano, Oh et al. 2020, Addiction 2024) Despite the significant personal, societal and financial burdens imposed by cocaine use, there are currently no FDA-approved treatments. Addressing gaps in our knowledge about the underlying mechanisms that lead to CUD would aid in identifying novel targets and developing effective treatment and preventative strategies.

Among addictive substances, cocaine has one of the highest risks of addiction (Goldstein and Kalant 1990). Twin studies indicate that the heritability (*h^2^*) of cocaine dependence ranges from 0.4 - 0.7 indicating that genetics play a significant role (Goldman, Oroszi et al. 2005, Ducci and Goldman 2012). Human GWAS for cocaine dependence identified a significant association with a single nucleotide polymorphism (SNP) in the FAM53B (family with sequence similarity 53, member B) gene on chromosome (Chr) 10 (Gelernter, Sherva et al. 2014) in a region that was identified by linkage analysis in a previous study (Gelernter, Panhuysen et al. 2005). Despite a few successful studies, GWAS for CUD have been limited by insufficient sample sizes that reduce power to detect the likely numerous genetic variants that drive cocaine use and CUD (Goldman, Oroszi et al. 2005).

Genome-wide mapping approaches have been employed in mice and overcome some of the obstacles present in humans. The use of experimental mice allows for control over drug exposures, minimizes environmental variability and provides access to brain tissue for mechanistic studies. Moreover, gene by drug interactions can be assessed by studying the effects of drug exposure on different genetic backgrounds. Genetic mapping in mice has identified numerous quantitative trait loci (QTL) for addiction-relevant behaviors (Phillips, Huson et al. 1998, Dickson, Miller et al. 2016, Bubier, Philip et al. 2020, Vadasz and Gyetvai 2020, Bagley, Khan et al. 2022). In rodents, acute psychomotor response to cocaine is commonly used as a measure of initial sensitivity to the drug. Human studies have shown that initial response to a drug predicts future use and abuse (de Wit and Phillips 2012). QTL studies for initial cocaine sensitivity have been conducted using standard recombinant inbred (RI) strains (Tolliver, Belknap et al. 1994, Miner and Marley 1995, Phillips, Huson et al. 1998, Jones, Tarantino et al. 1999, Boyle and Gill 2001, Gill and Boyle 2003, Gill and Boyle 2008, Boyle and Gill 2009) or C57BL/6 substrains (Kumar, Kim et al. 2013) and have identified numerous genomic regions. To date, however, only one genetic variant in the *Cyfip2* gene has been identified and validated for initial locomotor sensitivity (Kumar, Kim et al. 2013). Identification of genes that influence initial locomotor sensitivity to cocaine will inform mechanistic studies that will further our understanding of specific factors that might drive risk for CUD.

In the present study, we utilized a new resource, the Collaborative Cross (CC). The CC was created by crossing eight inbred mouse strains including five classical strains – A/J (A), C57BL/6J (B6J), 129S1/SvImJ (129), NOD/ShiLtJ (NOD), NZO/HlLt (NZO) and three wild-derived strains – WSB/EiJ (WSB), CAST/EiJ (CAST), PWK/PhJ (PWK). The CC is unique in its genetic diversity as the eight inbred founders represent the three subspecies of *Mus musculus* (Churchill, Airey et al. 2004, Srivastava, Morgan et al. 2017). The extensive genetic diversity allows for novel combinations of alleles and the observation of expanded phenotypic diversity in comparison to traditional inbred strains and RI panels (Rasmussen, Okumura et al. 2014, Graham, Thomas et al. 2015, Gralinski, Ferris et al. 2015, Mosedale, Kim et al. 2017). Moreover, the availability of an expanded set of genetic and genomic tools developed to support CC studies allows for systems genetic analysis and the ability to move more quickly from QTL to candidate gene analyses (Ball, Bogue et al. 2024).

In a previous study, we identified two CC strains that exhibit low (CC041/TauUnc) or high (CC004/TauUnc) initial locomotor response to cocaine as well as differences in their ability to acquire intravenous cocaine self-administration (Schoenrock, Kumar et al. 2020). We crossed both of these CC strains to C57BL/6NJ (B6N) to generate two separate F2 mapping populations. QTL mapping using the CC041 X B6N F2 identified genome-wide significant QTLs on Chrs 7, 11 and 14. A similar mapping strategy in the CC004 X B6N F2 yielded only one suggestive locus. We used extensive *in silico* bioinformatic analyses to select candidate genes at the CC041 Chr 7 and 11 QTL peaks. Candidate genes with a high likelihood of being causal can be further characterized to test their role in addiction-relevant behaviors.

## METHODS AND MATERIALS

### Animals

Mice from the CC004/TauUncJ (CC004) strain were purchased from the Jackson Laboratory in November 2015. Mice from the CC041/TauUnc (CC041) strain were purchased from the Systems Genetics Core Facility at the University of North Carolina (UNC; http://csbio.unc.edu/CCstatus/index.py) in January 2016. CC004 and CC041 mice were reciprocally bred to C57BL/6NJ (B6N; RRID:MGI:3056279) mice purchased from The Jackson Laboratory (Bar Harbor, ME) to generate CCxB6N and B6NxCC F1s (denoted as BCF1 and CBF1). These F1s were crossed in all combinations (CCxB6N X CCxB6N, CCxB6N X B6NxCC, B6NxCC X CCxB6N, B6NxCC X B6NxCC denoted as BCBC, BCCB, CBCB and CBBC) to generate each of the two (CC004 and CC041) F2 populations.

Animals were housed in a specific pathogen-free facility on a 12-hour light/dark cycle with lights on at 7:00 A.M. Food and water were available *ad libitum* throughout the experiment. Breeder diet was Harlan Teklad 2919 and post-weaning diet was Harlan Teklad 2920 (Envigo, Frederick, MD, USA). All procedures were approved by the UNC Institutional Animal Care and Use Committee and followed guidelines set forth by the National Institutes of Health Guide for the Care and Use of Laboratory Animals.

### Drugs

Cocaine HCl (Sigma-Aldrich, St. Louis, MO) was dissolved in 0.9% saline and a dose of 20 mg/kg of body weight was used to test initial locomotor response in a 3-day open field (OF) test.

### Phenotyping for initial locomotor sensitivity to cocaine

Founder strain (CC004 and CC041), F1 and F2 mice were tested for initial locomotor response to cocaine using a three-day open field (OF) assay. Total numbers of mice tested for each group are shown in Table 1. On Days 1 and 2, mice were administered saline at a volume of 0.1ml/g via the intraperitoneal (i.p.) route and immediately placed into a 43.2 x 43.2 x 33 cm OF arena (ENV-515-16, Med Associates, St. Albans, VT, USA). Mice were tracked for the entire session by infrared detectors that surrounded the arena at 2.54 cm intervals on the x, y, and z axes. After 30 minutes, mice were removed from the OF and returned to the home cage. On Day 3, mice were given an i.p. injection of 20mg/kg cocaine at a volume of 0.1ml/g and placed in the OF for 30 minutes. On each day, distance moved (in centimeters) was recorded using behavioral software provided by the OF manufacturer (Med Associates; RRID:SCR_014296). Total distance moved during the entire 30-minute assay was used as a dependent variable.

**Table 1:** Number of mice phenotyped in CC004 and CC041 mapping populations.

| Population | Generation | Cross | # F | # M | Total |
| --- | --- | --- | --- | --- | --- |
| Cocaine low response | F0 | CC041/TauUnc (CC041) | 12 | 12 | 24 |
|  |  | C57BL/6NJ (B6N)* | 11 | 13 | 24 |
|  | F1 | B6NxCC041 | 13 | 13 | 26 |
|  |  | CC041xB6N | 11 | 12 | 23 |
|  |  |  |  |  | **49** |
|  | F2 | B6NxCC041 X B6NxCC041 | 58 | 62 | 120 |
|  |  | CC041xB6N X CC041xB6N | 59 | 53 | 112 |
|  |  | CC041xB6N X B6NxCC041 | 66 | 63 | 129 |
|  |  | B6NxCC041 X CC041xB6N | 37 | 50 | 87 |
|  |  |  |  |  | **448** |
| Cocaine high response | F0 | CC004/TauUnc (CC004) | 11 | 14 | 25 |
|  |  | C57BL/6NJ (B6N)* | 11 | 13 | 24 |
|  | F1 | B6NxCC004 | 9 | 8 | 17 |
|  |  | CC004xB6N | 9 | 10 | 19 |
|  |  |  |  |  | **36** |
|  | F2 | B6NxCC004 X B6NxCC004 | 38 | 39 | 77 |
|  |  | CC004xB6N X B6NxCC004 | 38 | 39 | 77 |
|  |  | CC004xB6N X CC004xB6N | 38 | 40 | 78 |
|  |  | B6NxCC004 X CC004xB6N | 39 | 38 | 77 |
|  |  |  |  |  | **309** |

\*Same animals

### Statistical analyses

An ANOVA of the effects of day (1, 2, 3), strain (CC004, CC041, F1 and F2) and sex on distance moved in the OF was performed using SPSS v28 for Mac OS (IBM Corp; RRID:SCR_016479). A separate ANOVA tested the effects of strain and sex on locomotor response to cocaine on Day 3 minus baseline activity on Day 2 (Day 3 – Day 2) was also conducted. For both analyses, mice from reciprocal crosses in the F1 and F2 were analyzed as separate groups to identify parent of origin effects that might impact mapping analyses. *Post hoc* Tukey’s *HSD* was used to follow-up on any significant main effects of strain. Graphs were generated using Graphpad Prism 10 for Mac OS (GraphPad Software, LLC; RRID:SCR_002798).

### Genotyping

Genotyping was performed for the 444 CC041 X B6N F2 mice and 151 CC004 X B6N F2 mice. A total of 309 CC004 X B6N F2 mice were produced and behaviorally characterized, but only the phenotypic extremes were selected for genotyping. Mice were euthanized immediately following testing on Day 3 and DNA was extracted from tail tissue of CC parents, F1 breeders, representative B6N samples and the F2 population using the DNeasy Blood & Tissue kit (Qiagen). Genotypes for the CC041 X B6N F2s were determined using the Mouse Universal Genotyping Array (MUGA) that consists of 7851 SNP markers on an Illumina Infinium platform that are distributed throughout the genome (average spacing of 325 kb) and were chosen to be maximally informative for the eight founder strains of the CC (Morgan, Fu et al. 2015). Genotypes for the CC004 X B6N F2s were determined using the fourth iteration of the Mouse Universal Genotyping Array (MiniMUGA). The MiniMUGA contains a total of 10,821 genomic probes that are most informative SNPs from 120 classical inbred lines of mice (Sigmon, Blanchard et al. 2020, Blanchard, Sigmon et al. 2024). Nucleotide genotypes were processed and converted to haplotype calls (i.e. B6N, CC, or HET) for use in QTL mapping using the *argyle* package (version 0.2.0) in R Studio (Morgan 2015). A series of genotype checks were performed, and markers were eliminated for any of the following reasons: markers were not informative between the CC and B6N, did not meet the Chi-square distribution of expected genotypes for an intercross population (showed segregation distortion) or had a large number of missing calls (>40). In the CC041 X B6N F2 population, this strategy resulted in a final marker set that included 2701 markers with an approximate average spacing of 1 megabase (Mb) and maximum gap of 18 Mb. For the CC004 X B6N F2 population, the final marker set included 2394 markers with an average spacing of approximately 2 Mb and a maximum gap of 72 Mb on Chr 15. Both marker sets have more than ample coverage for an F2 population.

### QTL mapping

All data were transformed using a two-step inverse rank transformation to normality in SPSS. QTL mapping was performed using the *qtl* package (version 1.40-8;(Broman, Wu et al. 2003, Broman 2014) in R Studio (version 1.0.136). Single scan QTL analyses using the *scanone* function were performed for transformed total distance data for Day 1, Day 2 and Day 3 and D3-D2 distance. Sex and F2 breeding cross direction were included as covariates. A Haley-knot regression approximation model of interval mapping was used based on the density of the genotyping array and the amount of recombination present in the F2 intercross. Genome-wide significant thresholds for logarithm of the odds (LOD) scores (measure of genotype to phenotype association) were determined using 1000 permutations. For each QTL peak identified, the 1.5 LOD support interval was used as it can be translated to having a 95% likelihood for containing the casual variant (Dupuis and Siegmund 1999). The MUGA or MiniMUGA markers closest to the outer limits of the 1.5 LOD interval were identified and used as the megabase (Mb) locations flanking the region (mm10, GRCm38). Genotype data provided in Srivastava, Morgan et al. (2017) was used to determine the CC parental strain haplotype in the 1.5 LOD intervals for each QTL. Using the *fitqtl* function and an equation assuming an additive model, (e.g. *y = QTL 1 + QTL 2 + QTL 3*), the amount of variance explained for the QTL peaks of a given phenotype was estimated.

Two QTL analysis using the *scantwo* function in *R/qtl* was performed for the significant QTLs identified for each of the measures with sex and cross direction as covariates. Both full (LOD*_fv1_*) and additive models (LOD*_av1_*), allowing for the possibility of epistasis or not, respectively, were fit by comparing pairs of loci on the two chromosomes to the single-QTL model. LOD*_i_* indicates evidence for an interaction of the two loci by comparing the fit of the two models (LOD*_fv1_* -LOD*_av1_*). Thresholds of 6, 4, 3 for LOD*_fv1_*, *_i_*, *_av1_* were used to determine significant pairs of QTLs (Broman and Sen 2009). Effect plots at each QTL peak and for significantly interacting markers were generated using the *effectplot* function.

### Prioritizing candidate genes in CC041 QTL regions

We prioritized potential candidate genes in the QTL intervals using a multi-step process (**Fig 4**). We queried the GenomeMUSter website (Ball, Bogue et al. 2024) (https://mpd.jax.org/genotypes) to identify SNPs that varied between CC041 and B6N in the Chr 7 (27.5-37.9Mb) and Chr 11 (12.6-37.7Mb) QTL intervals. We prioritized deleterious polymorphisms such as STOP loss or gain mutations as well as missense or non-synonymous amino acid mutations. In order to determine if amino acid changes were predicted to be tolerated or disruptive, the SNP ids were tested using SIFT (https://sift.bii.a-star.edu.sg/; RRID:SCR_012813)(Sim, Kumar et al. 2012). We also selected SNPs that reside in evolutionarily conserved residues by performing multi-species protein sequence alignment, retrieving protein amino acid sequences from UniProt (http://www.uniprot.org; RRID:SCR_002380)(UniProt 2023) and performing multiple sequence alignments using CLUSTAL Omega (https://www.ebi.ac.uk/Tools/msa/clustalo/ RRID:SCR_001591)(Sievers, Wilm et al. 2011).

## RESULTS

### Locomotor response to cocaine in the CC041 population

The number of mice tested in each mapping population is provided in **Table 1**. We performed an analysis of variance (ANOVA) of locomotor activity with day, strain and sex as independent variables. We observed that locomotor activity decreased significantly on day 2 compared to day 1 (*p* = 3.0 x 10^-6^) and increased significantly on day 3 compared to both day 2 (*p* = 5.1 x 10^-9^) and day 1 (*p* = 5.1 x 10^-9^)**(Fig 1A)**. We observed a main effect of strain; C57BL/6NJ were significantly more active than CC041, F1 and F2 mice (all *p* < 0.001; **Fig 1B**) regardless of day. No parent of origin effect was observed in the F1 as BCF1 mice did not differ significantly from CBF1 mice (*p* > 0.05). In the F2, BCBC mice (N = 120) were significantly less active than BCCB mice (N = 87; *p* = 0.016). No other significant differences were observed among the different F2 cross types. Both BC and CB F1 mice showed a response similar to CC041, indicating that the low-responding phenotype is dominant. The CC041 X B6N F2 population showed a wide range of responses, although a large proportion of mice exhibited locomotor activity at the lower end of the phenotypic distribution appearing more similar to the CC041 parent, which is consistent with our observations in the F1 and parental strains (**Fig 1C**). We also observed significant strain by day (F_(14,1587)_ = 11.2; *p* = 6.5 x 10^-25^) and strain by sex (F_(7,1587)_ = 7.9; *p* = 3.8 x 10^-4^) interactions. Strain by day interactions generally reflected differences between strains in response to cocaine (or lack thereof) on day 3. Neither CC041 nor CB F1 mice differ in locomotor activity across days in comparison with all other strains and cross-types. Female mice of all F2 cross types had significantly higher locomotor activity than male mice (all *p* < 0.01) but the same sex difference was not observed in either F1 cross or the parent inbred strains.

**Figure 1.**
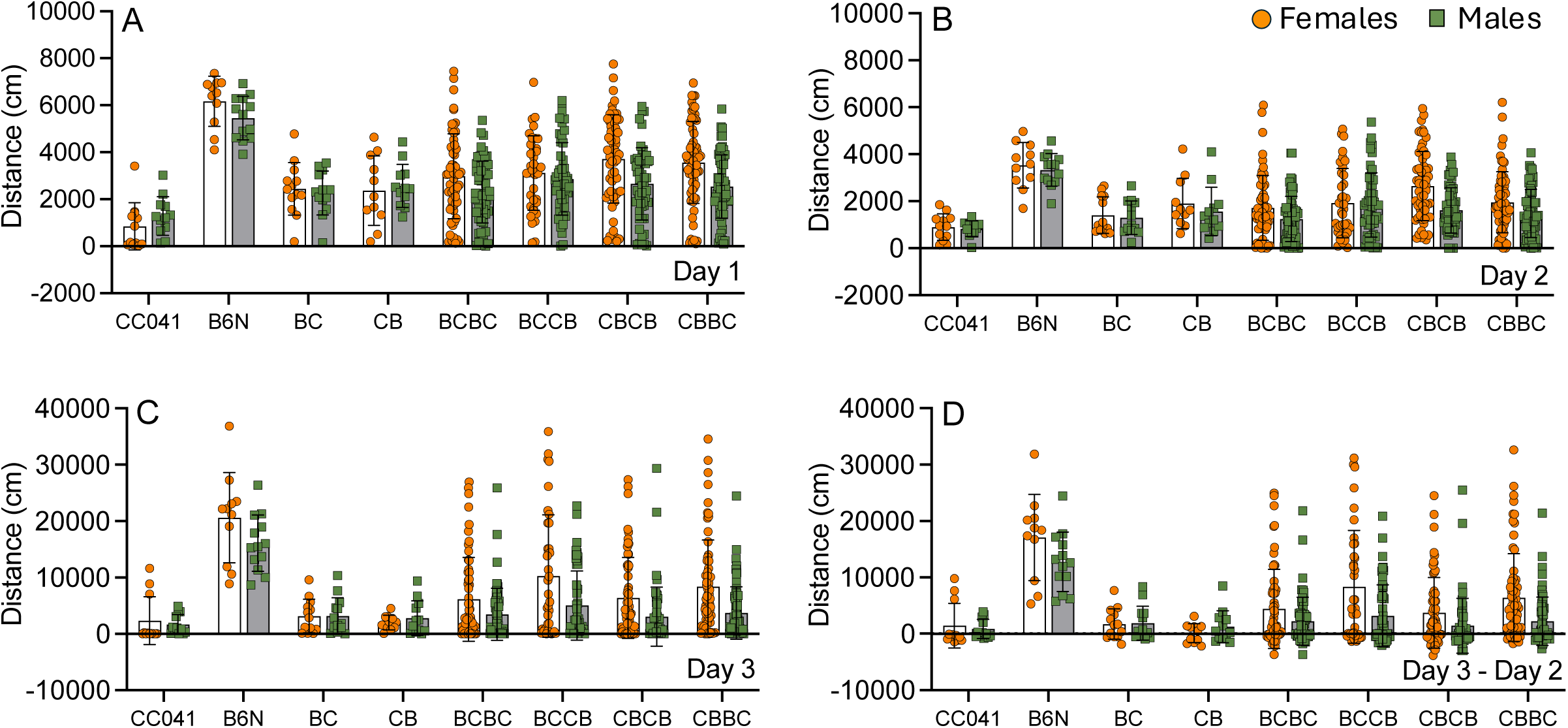
Locomotor activity in response to saline on Day 1 (A) and Day 2 (B), after cocaine exposure on Day 3 (C) and Day 3 minus Day 2 (D) in CC041 and B6N parent strains, reciprocal F1 mice and all F2 cross types. BC = B6N x CC041 F1, CB = CC041 x B6N F1, BCBC = BC F1 x BC F1, BCCB = BC F1 x CB F1, CBCB = CB F1 x CB F1 and CBBC = CB F1 x BC F1. Strain background of dam is listed first.

ANOVA of locomotor response to cocaine on Day 3 minus Day 2 yielded a significant effect of strain (F_(7,529)_ = 15.8; *p* = 7.1 x 10^-19^) and sex (F_(1,529)_ = 10.4; *p* = 0,001) but no strain by sex interaction (F_(7,529)_ = 1.5; *p* = 0.15). B6N mice have a significantly higher locomotor response to cocaine than all other strains (**Fig 1D**; *p* values all <1.0 x 10^-10^). Female mice exhibit a higher locomotor response to cocaine than male mice.

### Locomotor response to cocaine in the CC004 population

An ANOVA of locomotor activity with day, strain and sex as independent variables yielded significant main effects of day (F_(2,1200)_ = 585.3; *p* = 4.0 x 10^-178^), strain (F_(7,1200)_ = 15.8; *p* = 6.5 x 10^-20^) and sex (F_(1,1200)_ = 46.9; *p* = 1.2 x 10^-11^). As in our other population, activity decreased from day 1 to day 2 and increased significantly on day 3 (all p < 0.001; **Fig 2A-C**). CC004 mice had significantly higher activity than B6N, as well as the F1 and F2 crosses (all p < 0.01). BCF1 mice were significantly more active than CBF1 mice suggesting a parent of origin effect on the phenotype. No differences in locomotor activity were observed among the different F2 cross types. We also observed significant strain by day (F_(14,1200)_ = 7.9; *p* = 3.0 x 10^-16^), strain by sex (F_(7,1200)_ = 2.1; *p* = 0.037) and sex by day (F_(2,1200)_ = 23.0; *p* = 1.6 x 10^-10^) interactions.

**Figure 2.**
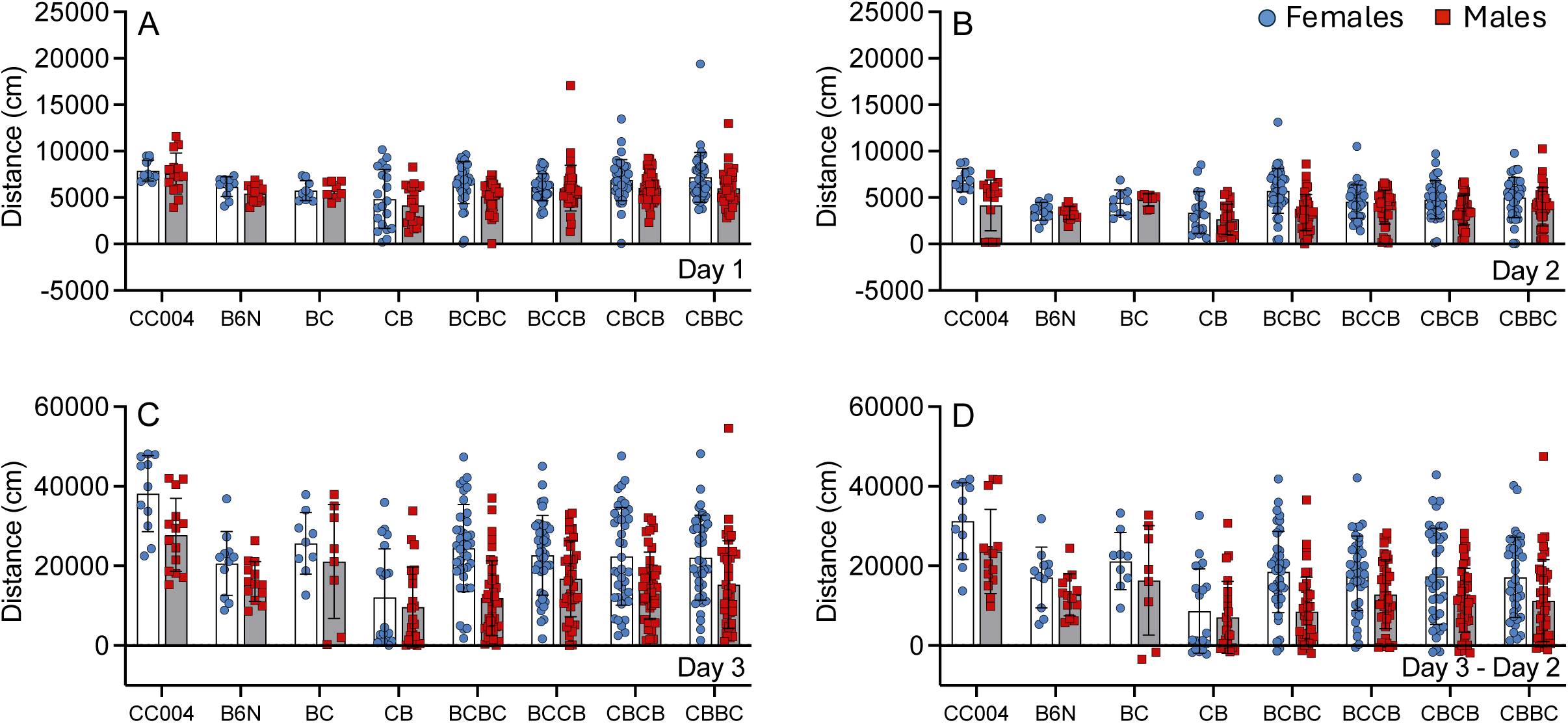
Locomotor activity in response to saline on Day 1 **(A)** and Day 2 **(B)**, after cocaine exposure on Day 3 (C) and Day 3 minus Day 2 (D) in CC004 and B6N parent strains, reciprocal F1 mice and all F2 cross types. BC = B6N x CC004 F1, CB = CC004 x B6N F1, BCBC = BC F1 x BC F1, BCCB = BC F1 x CB F1, CBCB = CB F1 x CB F1 and CBBC = CB F1 x BC F1. Strain background of dam is listed first.

ANOVA of locomotor response to cocaine on Day 3 minus Day 2 yielded significant strain (F_(7,401)_ = 9.9; *p* = 2.3 x 10^-11^) and sex (F_(1,401)_ = 25.6; *p* = 6.3 x 10^-7^) effects but no strain by sex interaction (F_(7,401)_ = 0.85; *p* = 0.54). CC004 mice are significantly more active in response to cocaine than all other strains (**Fig 2D**; all *p* < 0.001) except for BC F1 mice (*p* = 0.139)

### QTL on Chr 7, 11 and 14 for low initial locomotor response to cocaine in CC041 X B6N population

We conducted genetic mapping in our CC041 x B6N population and identified 3 significant QTL for locomotor response to cocaine (Day3-Day2 distance) on Chr 7, 11 and 14 (**Fig 3A**). QTLs that passed the p<0.1 genome-wide LOD threshold based on 1000 permutations are shown in **Table 2**. The QTL on Chr 7 had a peak at 17.2cM or 29.74Mb (LOD=4.21, *p*=0.023) and accounted for 4.3% of the variance. The effect plot at this peak showed that both the CC041/CC041 and HET genotypes were driving the low cocaine response, again consistent with our initial observations in the parental and F1 populations (**Fig 3B**). The LOD interval (27.5-40.3 Mb) of the Chr 7 peak is 12.8 Mb and contains 203 protein-coding genes. CC041 has a NOD haplotype that transitions to NZO at approximately 36.05Mb in this interval (**Table 2**). The QTL on Chr 11 had a peak at 14.3cM or 20.98Mb (LOD=6.54, *p*=0.001) and accounted for 6.1% of the variance. The effect plot at this peak showed that the B6N/B6N genotype was driving the low cocaine response (**Fig 3C**) indicating that this is a transgressive QTL. The LOD interval (12.6-37.7Mb) of the Chr 11 peak is 25.1 Mb and contains 106 protein-coding genes. CC041 has a NOD haplotype that transitions to a WSB haplotype at approximately 36.8 Mb. The Chr 14 QTL had a peak at 42.7cM or 79.67Mb (LOD=3.88, *p*=0.051) that accounted for 4% of the variance. The effect plot at this peak showed that the CC041/CC041 genotype was driving the low cocaine response (**Fig 3D**). The LOD interval (9.1-97.4) on Chr 14 was very large spanning most of the chromosome (88.3 Mb) due to the appearance of multiple peaks towards the proximal end, therefore we did not perform follow-up analysis of potential candidate genes in this region.

**Figure 3.**
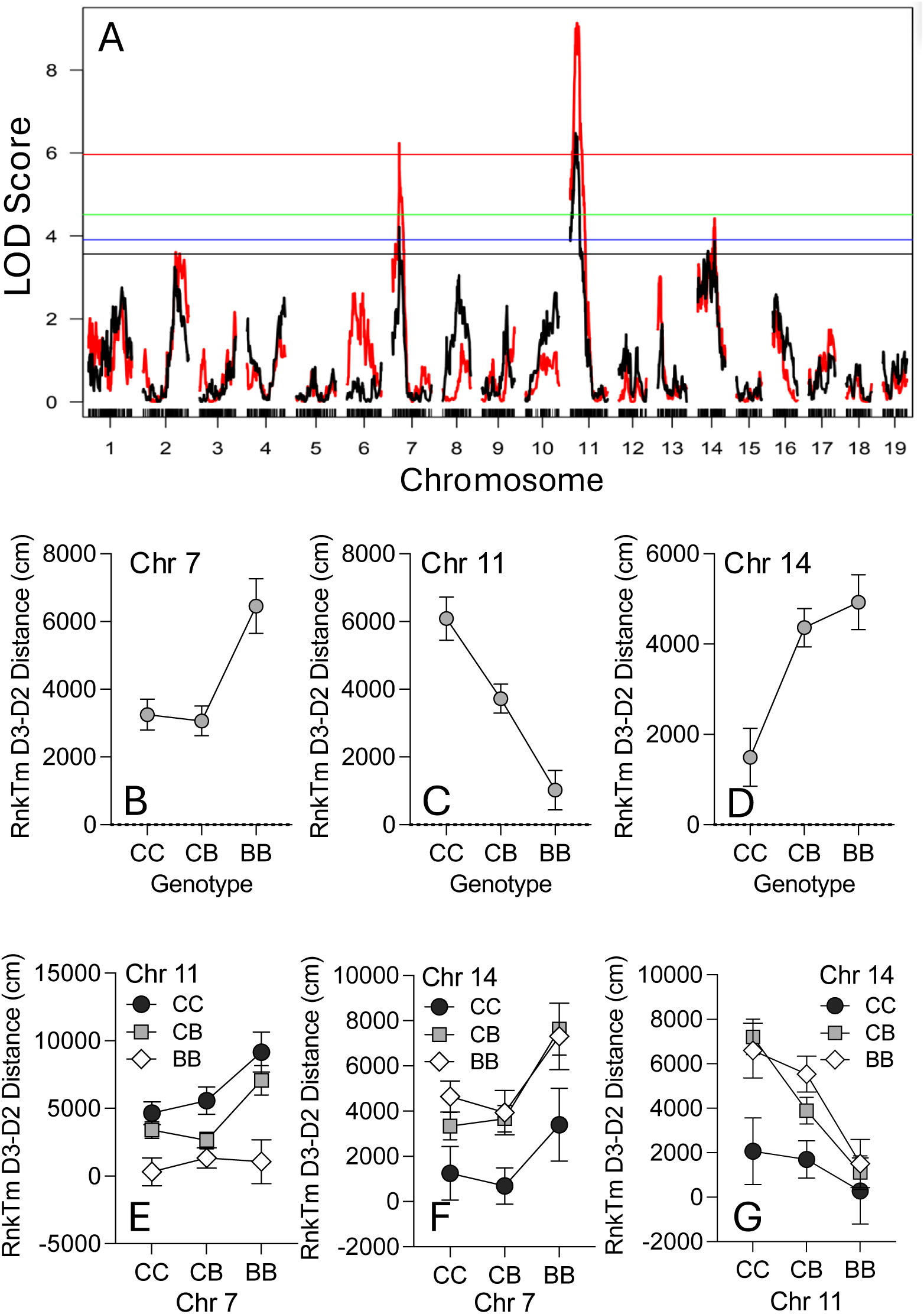
**(A)** Single scan QTL for initial cocaine response measured as Day 3 minus Day 2 (red) compared to Day 2 locomotor activity (black). Genome-wide significant LOD thresholds based on 1000 permutations for *p*=0.001 (red line), *p*=0.01 (green line), *p*=0.05 (blue line), and suggestive at *p*=0.1 (black line). QTL effect plots for Chr 7 **(B)**, 11 **(C)** and 14 **(D)** and two-scan interaction plots for Chr 7 vs Chr 11 **(E)**, Chr 7 vs Chr 14 **(F)** and Chr 11 vs Chr 14 **(G)**. CC = CC041/CC041 homozygote, CB = CC041/B6N heterozygote and BB = B6N/B6N homozygote. Error bars are SEM.

**Table 2.** QTL regions identified in the 3-day open field test.

|  |  | Chr | Mb | Marker Name | LOD | p-value | 1.5 LOD Interval (Mb) | Interval Size (Mb) | Decreasing Allele | CC Parent Haplotype |
| --- | --- | --- | --- | --- | --- | --- | --- | --- | --- | --- |
| CC041 X B6N F2 | Day 1 Distance | 6 | 97.0 | backupUNC060149691 | 6.29 | 0.001 | 51.5 - 111.4 | 59.9 | CC041 | CC |
|  |  | 11 | 35.7 | UNC111423133 | 3.88 | 0.055 | 18.0 - 70.8 | 52.8 | B6N | DD/HH |
|  |  | 14 | 84.6 | JAX00385742 | 6.92 | 0.001 | 73.3 - 94.7 | 21.4 | CC041 | FF |
|  | Day 2 Distance | 6 | 15.2 | UNC060057271 | 5.33 | 0.004 | 21.9 - 114.0 | 92.1 | CC041 | CC |
|  |  | 7 | 28.9 | UNC070790469 | 3.76 | 0.070 | 4.0 - 41.5 | 37.5 | CC041 | DD/EE/DD |
|  |  | 11 | 35.7 | UNC111423133 | 5.00 | 0.004 | 21.7 - 51.6 | 29.9 | B6N | DD/HH |
|  | Day 3 Distance | 2 | 152.7 | UNC020230950 | 3.62 | 0.089 | 141.5 - 180.8 | 39.3 | CC041 | HH/CC/EE |
|  |  | 7 | 29.7 | UNC070628845 | 6.24 | 0.001 | 27.5 - 37.9 | 10.4 | CC041 | DD/EE |
|  |  | 11 | 30.9 | backupJAX00025840 | 9.14 | 0.000 | 18.0 - 38.6 | 20.6 | B6N | DD/HH |
|  |  | 14 | 79.7 | backupUNC140618860 | 4.43 | 0.013 | 12.6 - 97.2 | 84.6 | CC041 | DD/CC/FF |
|  | Day 3-Day 2 Distance | 7 | 29.7 | UNC070628845 | 4.22 | 0.023 | 27.5 - 37.9 | 12.8 | CC041 | DD/EE |
|  |  | 11 | 20.9 | UNC110035818 | 6.48 | 0.001 | 12.6 - 37.7 | 25.1 | B6N | DD/HH |
|  |  | 14 | 79.7 | backupUNC140618860 | 3.88 | 0.056 | 9.1 - 97.5 | 88.3 | CC041 | DD/CC/FF |
| CC004 X B6N F2 | Day 3-Day 2 Distance | 11 | 118.1 | mUNC20502282 | 3.76 | ns | 4.9 - 118.5 | 113.6 | B6N/CC041 | AA/CC/DD |

All QTL regions above the suggestive genome-wide threshold ( $p=0.01$ ) using 1000 permutation for phenotypes in the 3-day open field test in the low-cocaine responding F2 population. Decreasing allele indicates the genotype associated with the lowest phenotype (CC041 = CC041/TauUnc; B6N = C57BL/6NJ). Collaborative Cross (CC) parental strain haplotype in QTL region is indicated by CC (129S1/SvImJ), DD (NOD/ShiLtJ), EE (NZO/HILtJ), FF (CAST/EiJ), HH (WSB/EiJ). Chr = chromosome; cM = centimorgan; Mb = megabase position in mm10 mouse genome

Single QTL analysis of Day 1, Day 2 and Day 3 distance revealed several overlapping regions (**Table 2, Suppl Fig 1**). Interestingly, a peak on Chr 11 was identified for all days in similar locations to the peak identified for cocaine locomotor response (peaks at 30.88 or 35.7 Mb compared to 20.9 Mb) and in every case, the B6N/B6N genotype was associated with the low response. A QTL peak on Chr 7 was identified for Day 2 and Day 3 distance with the Day 3 peak being the same as the one identified for Day 3-Day 2 and in every case, the CC041/CC041 and CC041/B6N genotypes was associated with low response. A QTL peak on Chr 14 was identified for Day 1 and Day 3 with the Day 3 peak being the same as the one identified for Day 3-Day 2 and in all cases only the CC041/CC041 genotype showed the low response. Two separate QTLs on Chr 6 were specific to non-cocaine (saline) exposures on Day 1 (peak at 97.0 Mb) and Day 2 (peak at 37.0 Mb). For both Chr 6 QTL, the CC041/CC041 genotype was associated with low locomotor response. A QTL on Chr 2 (peak at 152.7 Mb) was unique for the Day 3 distance phenotype with the CC041/CC041 genotype driving the low response.

Collectively, the single scan QTL analysis in the low responding population indicates that QTL identified are associated with locomotor phenotypes on multiple days in the 3-day OF test, a finding that could be due to the significant correlation among the phenotypes (**Suppl Table 1**). Pearson correlation showed that Day 1, Day 2, Day 3 and Day 3-Day 2 distance were all significantly correlated (r(6)=0.31-0.99; *p≤*1×10^-15^). The strongest correlation was observed between Day 3 distance and D3-D2 distance (r(6)=0.99; *p≤*1×10^-15^).

### Two-locus analysis shows additive effect of QTLs for low COC response in CC041 X B6N population

Most complex traits, such as initial drug sensitivity, are thought to be due to the action of multiple genetic loci, some of which may act in concert to affect the phenotype. Therefore, we performed a two-locus analysis to determine if the QTLs identified in the single scan are interacting and if so, in what manner (i.e. additive or epistatic). For each phenotype, only the chromosomes with significant QTLs in the single scan were analyzed. **Table 3** shows all loci that met the threshold in either a full model (allows for possibility of epistasis; LOD*_fv1_* ≥ 6.0) or additive model (assumed no epistasis; LOD*_av1_* ≥ 3.0).

**Table 3.**
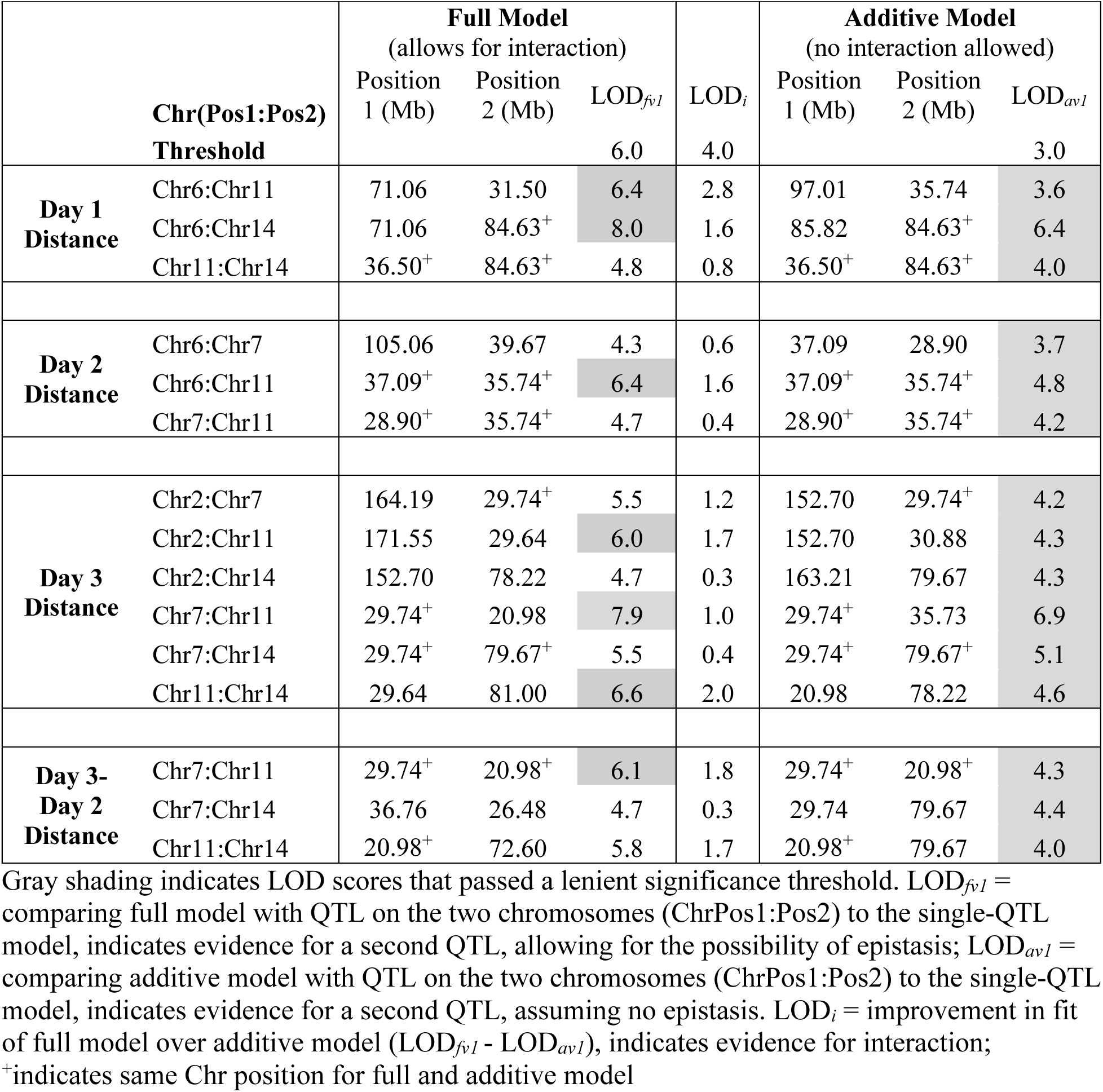
Two scan QTL results for CC004 X B6N F2 mapping population.

For Day3-Day2, we found strong evidence for an interaction among a pair of QTL at the same locations identified in the single scan analyses on Chrs 7 and 11. The presence of a homozygous B6N genotype on Chr 11 results in low response to cocaine regardless of the genotype on Chr 7 (**Fig 3E**). There was also evidence for an interaction between QTL on Chrs 7 and 14. Although the Chr 7 QTL effect is preserved across genotypes, CC041 homozygosity at the Chr 14 locus results in reduced locomotor response to cocaine across all three Chr 11 genotypes. There was also evidence for interaction between a pair of QTL on Chrs 11 and 14 (**Fig 3F**). CC041 homozygosity at the Chr 14 locus represses the allelic effects of the Chr 11 QTL on locomotor response to cocaine (**Fig 3G**).

### QTL in CC004 X B6N population

A genome-wide scan in the CC004 X B6N F2 cross identified a suggestive QTL peak at the distal end of Chr 11 at 118 Mb or (LOD = 3.76). (**Table 2**). No other QTL were identified in this cross.

### Prioritization of candidate genes within the Chr 7 and Chr 11 QTL regions identified in CC041 X B6N population

The strategy outlined in **Fig 4** was used to identify potential candidate genes for COC locomotor response at the QTL regions on Chr 7 and Chr 11. The 12.8 Mb QTL region on Chr 7 contained 203 protein-coding genes. We identified 63,686 SNPs that differed between B6N and CC041. This included 283 different genomic features (73 Gene Models, 5 Mir-RNA, 29 *Scgb*-family members, 23 *Zfp* family members, 21 *RIK* or contig clones). There were zero stop loss or gain mutations and 152 missense mutations. Due to the one-to-many mappings (a SNP can be in multiple open reading frames), 168 mutations were determined to be tolerant, 21 were determined to be deleterious and 33 were unannotated (**Table 4**). Of the 21 deleterious mutations, only 3 (*Ryr1*, *Ffar3* and *Lgi4*) were in protein residues that were conserved across species (mouse, human and at least one other mammal) suggesting that SNP had functional significance. We excluded *Nudt19*, *Nphs1* and *Cd22* genes as the protein residue at the location of the SNP is not conserved across species suggesting non-conservation of function. We also excluded the *Zfp* or *Scgb* family proteins due to the size of each gene family and their redundancy in the genome. A variant in *Wdr62* resulted in an A137T missense mutation in the WDR62 protein. However, while the reference strain, B6J has an A at that location, rat, human, pig and the NOD strain have a T.

**Figure 4.**
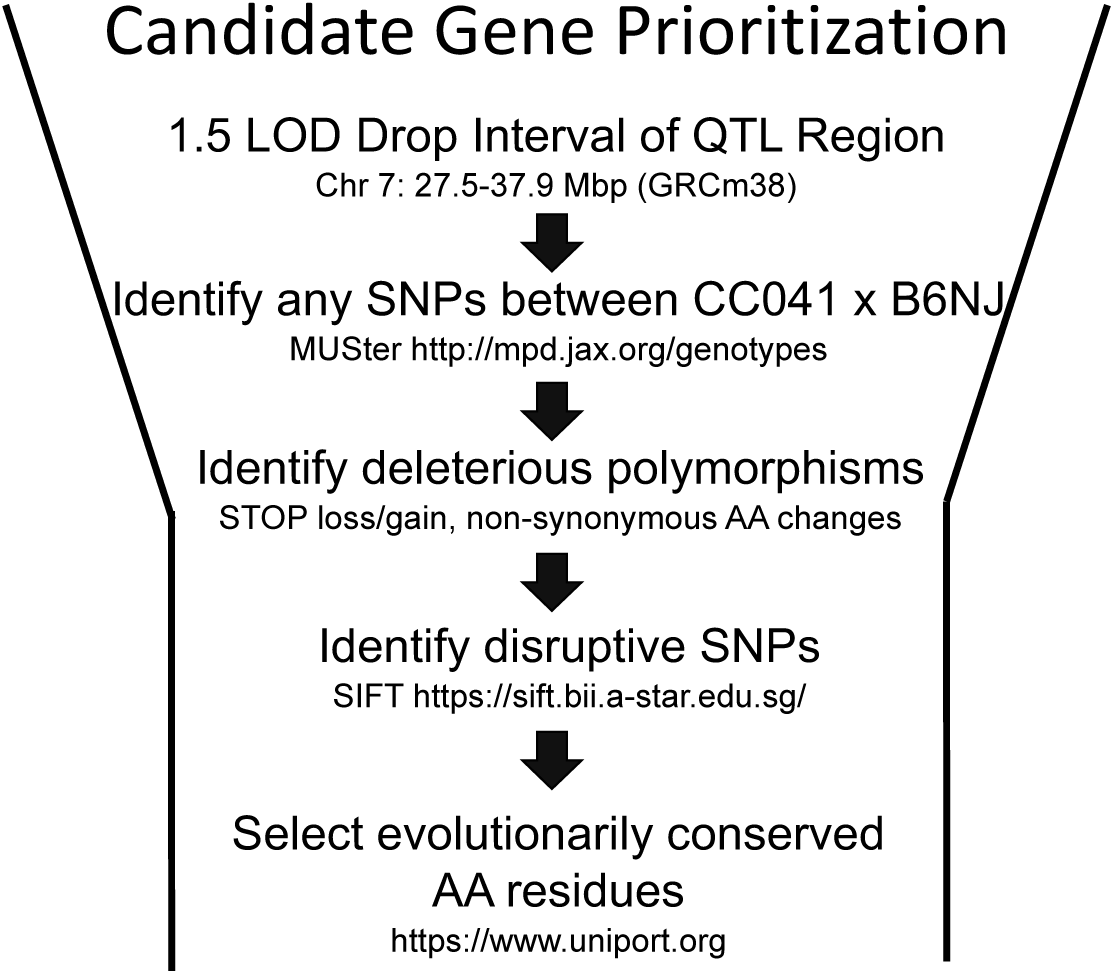
Strategy for identifying key candidate genes based on 1.5 LOD support interval and bioinformatic approaches to identify deleterious polymorphisms that are conserved and likely functionally relevant. Using the BioMart tool, we identified a list of protein-coding genes, lncRNA, and miRNAs. We eliminated any that were in a region of IBD between B6J and the CC parental strain using the Mouse Phylogeny Viewer. We also eliminated any genes not expressed in the brain. We prioritized genes that had a nonsynonymous SNP between B6J and the CC parental strain, specifically targeting SNPs in coding sequence, splice regions, stop regions, regulatory regions, frameshift and missense using the Sanger Institute SNP Viewer tool. Finally, we prioritized genes that met all three of the following criteria: previously identified with a phenotype of interest (i.e. brain structure/function or behavioral); expression differentially regulated by cocaine (using www.GeneWeaver.org); region that overlapped with previous QTL studies for initial cocaine-induced locomotion.

**Table 4.** Number and class of genetic variants between CC041 and B6N at Chr 7 and Chr 11 QTL intervals.

| Variation Type | QTL Interval |  |
| --- | --- | --- |
|  | Chr 7 | Chr 11 |
| 3' UTR | 486 | 810 |
| 5' UTR | 151 | 129 |
| Frameshift | 0 | 2 |
| Inframe insertion | 0 | 1 |
| Missense | 152 | 104 |
| Non-coding exon | 1152 | 1133 |
| Splice acceptor | 2 | 2 |
| Splice donor | 4 | 3 |
| Splice region | 79 | 99 |
| Synonymous | 238 | 281 |

The 25.1 Mb QTL region on Chr 11 contained 298,431 SNP differences between CC041and B6N. SNPs were located in 140 protein coding genes (18 Rik, 33 GeneModel) and 3 miRNA. Of those SNPs, 157,134 have registered rsID numbers. There were 658 coding non-synonymous substitutions (104 classified as missense variants), 281 synonymous substitutions, 268 lincRNA, 810 present in UTR 3’ and 129 in UTR5’ regions. 642 of the SNPs located in coding sequence were classified as tolerated. There are 16 deleterious SNPs that are located in 7 genes. Of those, only *Vrk2*, *Clhc1* and *Il9r* SNPs have a strong deleterious effect in a residue that is conserved across species. In addition, *Meis1* and *Psme4* are alternatively spliced in B6 vs CC041 and *Wdpcp* has an amino acid duplication in CC041 (rs230611425) that is not present in B6 and includes a conserved residue.

## DISCUSSION

We employed F2 intercrosses of a traditional inbred mouse strain (B6N) to two CC strains (CC041 & CC004) to identify genetic loci associated with locomotor sensitivity to cocaine. The CC004 x B6N cross identified only one suggestive locus, but we were able to map three significant cocaine-induced locomotor response QTLs on Chrs 7, 11 and 14 in the CC041 x B6N cross. The association between genotype and phenotype on Chr 7 matches the mode of inheritance observed in the parental lines. At the Chr 11 QTL, however, we observed reduced locomotor response to cocaine that was associated with the B6N genotype, contrary to the parental phenotypes. All three CC041 x B6N QTL appear to be acting in an additive manner, although there is evidence for an epistatic interaction based on allelic status at Chr11 with both the Chr 7 and 14 loci. Using a bioinformatic approach, we identified potential candidate genes in the Chr 7 and Chr 11 QTL regions that could be mediating cocaine locomotor response. Below we compare our QTL regions to those previously identified and discuss several high-priority candidate genes.

### Previous QTL studies for initial locomotor sensitivity to cocaine

Several previous studies have identified QTL for initial locomotor response to cocaine using panels of RI or congenic strains (Tolliver, Belknap et al. 1994, Miner and Marley 1995, Phillips, Huson et al. 1998, Jones, Tarantino et al. 1999, Boyle and Gill 2001, Gill and Boyle 2003, Gill and Boyle 2008, Boyle and Gill 2009). Three of these studies have identified regions that overlap with our Chr 7 and 11 QTL regions. Gill and Boyle (2003) used congenic strains from the AcB/BcA panels derived by backcrossing the A/J inbred strain to B6J or the B6J strain to A/J and identified a QTL associated with initial cocaine locomotor response on Chr 7 for which increased cocaine response was driven by the B6J donor allele. Using a 25 Mb region around the peak at 51.7 Mb yields a region from 33.2 to 76.9 Mb which overlaps with the proximal end of our Chr 7 QTL interval. Tolliver, Belknap et al. (1994) used C57BL/6J x DBA/2J (BXD) recombinant inbred strains to identify a locus on Chr 11 at 12.3 Mb where the B6J genotype was associated with higher cocaine response compared to DBA/2J. Using a 25 Mb region around the peak yields a QTL region of 0 to 33.4 Mb which overlaps with our Chr 11 QTL interval. Finally, Jones, Tarantino et al. (1999) used BXD strains to identify a QTL interval on Chr 11 at 12.3 – 19.7 Mb at which the B6J genotype was associated with increased cocaine response. This QTL interval overlaps with our Chr 11 region as well as the locus identified by Tolliver et al (1994).

Kumar, Kim et al. (2013) identified a QTL peak on Chr 11 (1.5 LOD interval 35-57 Mb) that is associated with locomotor response to cocaine in an F2 intercross of B6 substrains, B6N and B6J. A causal variant in the *Cyfip2* gene was associated with low response to cocaine in B6N mice. While *Cyfip2* does not fall within our Chr 11 QTL LOD support interval, we cannot rule out the possibility that the *Cyfip2* gene is causing the B6N-driven low cocaine response we observed.

CC004 and CC041 also differ in acquisition of cocaine self-administration (Schoenrock, Kumar et al. 2020). Dickson et al. (Dickson, Miller et al. (2016) identified QTL for cocaine self-administration in BXD mice on Chr 7 (30.4-30.6 Mb) within our Chr 7 QTL interval and on Chr 11 (46.18-50.57 Mb). The Chr 11 QTL is slightly outside our region but does contain the *Cyfip2* gene. For both QTL, mice with the B6J genotype showed higher and more consistent cocaine infusions at lower doses of cocaine compared to animals with the DBA genotype.

### Potential candidate genes at the Chr 7 locus controlling locomotor response to cocaine

Previous studies have used a variety of techniques to prioritize candidate genes including pre-existing knowledge about the function of genes in relation to the drug’s effects (i.e. genes involved in the dopaminergic system), genes with parental strain polymorphisms, and correlation with gene expression changes. We used a systematic approach (**Fig 4**) to prioritizing candidate genes based on the nature of identified SNPs. We prioritized the predicted impact of all SNPs, moving from most deleterious to least deleterious. For coding non-synonymous (missense) SNPs, we considered the resulting amino acid change and the conservation of the altered residue across species. Our approach resulted in the identification of three candidate genes within the Chr 7 interval, *Ryr1, Ffar3* and *Lgi4*. We briefly discuss evidence from the literature that lends support for each potential candidate. Further studies that consider the phenotypic consequences of disrupting gene function are necessary to demonstrate the veracity of each candidate.

The ryanodine receptor gene, *Ryr1*, has a SNP (rs47114500) that results in a G to A transition and a change from alanine to valine at residue 1118. The alanine at that residue is conserved in mouse, human and rabbit suggesting functional significance. Ryanodine receptors (*RyRs*) are a family of intracellular receptors responsible for Ca2+ release to support a number of intracellular actions (Chan, Mayne et al. 2000, Chen, Tang et al. 2008, Grosskreutz, Van Den Bosch et al. 2010, Zundorf and Reiser 2011, Forostyak, Forostyak et al. 2016, Kilpatrick, Magalhaes et al. 2016, Egorova and Bezprozvanny 2019, Karagas and Venkatachalam 2019). Previous studies have identified an increase in *Ryr1* gene expression and elevation of RYR1 protein levels upon exposure to both methamphetamine and cocaine (Kurokawa, Mizuno et al. 2010, Kurokawa, Mizuno et al. 2011) that is regulated by activation of dopamine D1 receptors (Kurokawa, Mizuno et al. 2010, Kurokawa, Mizuno et al. 2011). Moreover, an induced mutant containing a human polymorphism exhibits hypolocomotion (Zvaritch, Kraeva et al. 2009) suggesting that the *Ryr1* gene plays a role in locomotor behavior.

*Ffar3* has a SNP (rs46782561) between CC041 and B6NJ that changes a proline to a leucine at residue 192 and has been previously associated with cocaine-induced methylation (5hmC) (Feng, Shao et al. 2015). This specific residue has been predicted by AlphaFOld (Jumper, Evans et al. 2021) to have hydrogen bonding with Thr 196 in its tertiary structure, contributing to the predicted detrimental effect of the variant. *Ffar3* encodes free fatty acid receptor 3, also known as GPR41, and is a type of G protein-coupled receptor essential for metabolism and immunity. (Kimura, Ichimura et al. 2020). This receptor has been shown to regulate the sympathetic nervous system by its expression on neurons involved in GBetaGamma-PLCbeta-MAPK signaling (Kimura, Inoue et al. 2011).

The leucine-rich repeat LGI family, member 4 gene (*Lgi4*) has a histidine to tyrosine substitution at residue 531 that is predicted to have deleterious effects. The histidine at this residue is conserved across mouse, human and bovine LGI4 proteins. LGI proteins play a role in synaptic transmission and myelination in the nervous system (Kegel, Aunin et al. 2013). In humans, mutations in LGI1 have been implicated in autosomal dominant lateral temporal lobe epilepsy (Nobile, Michelucci et al. 2009). A spontaneous mutation in the *Lgi4* gene (*Lgi4^clp^*) was identified in the colony at the Jackson Laboratory (Henry, Eicher et al. 1991) and was subsequently identified as a 225-bp insertion that results in loss of function. Studies using *Lgi4^clp^* mice established the gene’s function in Schwann cell signaling in the peripheral nervous system (Bermingham, Shearin et al. 2006). *In situ* hybridization studies characterized expression of *Lgi4* in the adult mouse medial septum, bed nucleus of the stria terminalis, substantia nigra and ventral tegmental area – brain regions that have been associated with addiction-relevant behaviors in mice (Herranz-Perez, Olucha-Bordonau et al. 2010). A mouse knockout of *Lgi4* (*Lgi4^tm1.1(KOMP)Vlcg^*) is homozygous lethal but heterozygous female mice exhibit decreased locomotor activity in comparison to wildtype females suggesting a role for *Lgi4* in locomotor activity (https://mousephenotype.org).

### Potential candidate genes at the Chr 11 locus controlling locomotor response to cocaine

An unexpected finding was the transgressive QTL on Chr 11 in which reduced locomotor response to cocaine was driven by the B6N allele rather than the CC041 allele. Initially we found this result somewhat surprising given that CC041 mice show lower cocaine response than B6N mice (see **Fig 1d**). We had crossed CC041 to B6N as B6N is highly genetically similar to the B6J CC founder strain, and as such would not introduce an entirely new genetic background while still maintaining mapping power. However, investigation of the founder contribution in CC041 revealed that B6J contributed only 0.05% to the genetic background. Consequently, we introduced allelic combinations that would not have been present in this CC strain making the transgressive nature of the Chr 11 QTL less surprising. Although causative variants in the Chr 11 region are likely not responsible for the phenotypic effect observed in CC041, they are clearly contributing to low response to cocaine and warrant further consideration. Our bioinformatic analysis yielded SNPs in three high-priority candidate genes on Chr 11, *Clhc1, Il9r* and *Vrk2*. Although *Clhc1* and *Il9r* are both expressed in the brain and have been implicated in neurobiological and behavioral phenotypes, *Vrk2* has the strongest evidence as a candidate gene.

A *Vrk2* SNP (rs13466583) in CC041 mice results in a proline to leucine substitution at residue 319, an important structural residue predicted to have deleterious effects due to its predicted role in hydrogen bonding (Varadi, Bertoni et al. 2024). The vaccinia-related kinase (VRK) family of proteins are members of the kinome, the complete set of protein kinases encoded by the genome (Caenepeel, Charydczak et al. 2004). *Vrk2* is a serine/threonine kinase that regulates transcription factors in pathways ranging from apoptosis to tumor growth. It has been identified as a gene of interest 185 times across 92 human GWAS studies (Sollis, Mosaku et al. 2023). Of relevance to substance use disorders, *Vrk2* was a human GWAS hit for opioid and cannabis use disorders (Xu, Toikumo et al. 2023), was associated 18 times each with schizophrenia and neuroticism measures, seven times with alcohol consumption or alcohol use disorder metrics (Karlsson Linner, Biroli et al. 2019, Kranzler, Zhou et al. 2019, Zhou, Sealock et al. 2020, Saunders, Wang et al. 2022, Kember, Vickers-Smith et al. 2023, Xu, Toikumo et al. 2023), and five times in smoking-related GWAS (Kichaev, Bhatia et al. 2019, Liu, Jiang et al. 2019, Xu, Li et al. 2020, Saunders, Wang et al. 2022, Xu, Toikumo et al. 2023). Variants in human VRK2 have also been associated with schizophrenia and major depressive disorder (Tesli, Wirgenes et al. 2016) (Li and Yue 2018) (Yin, Guo et al. 2023). *Vrk2* KO mice also display increased depressive-like behaviors, suggesting a role for the gene in the central nervous system.(Yin, Guo et al. 2023) Behavioral differences in these mice may be neurodevelopmental, as loss of *Vrk2* results in synaptic dysfunction and a reduction of dendritic spines in the ventral hippocampus (Yin, Guo et al. 2023).

We identified two deleterious polymorphisms in the coding sequence of the clathrin heavy chain linker domain containing 1 gene (*Clhc1*). A T to C transition changes a cysteine to arginine at residue 441 and an A to G transition changes threonine to alanine at residue 118. Polymorphisms in the human CLHC1 gene have been associated with bipolar disorder (Winham, Cuellar-Barboza et al. 2014), major depressive disorder (Coleman, Peyrot et al. 2020) and brain structure differences (van der Meer, Kaufmann et al. 2021). Behavior or brain structure phenotypes resulting from loss of function of *Clhc1* have not been reported in mice.

*Il9r* (interleukin 9 receptor) contains one SNP that differs between CC041 and B6N and results in a predicted deleterious coding-sequence substitution (S68L, S72L) in two of the five known *Il9r* transcripts. *Il9r* is differentially expressed in NA and hippocampus of High Responder (bHR) and Low Responder (bLR) rats (Clinton, Stead et al. 2011). An *Il9r* knock-out mouse has extensive defects related to regulatory T-cell development (Elyaman, Bradshaw et al. 2009) but no other behavioral phenotypes have been observed.

We also identified transcript variants in *Meis1* and *Psme4* genes. *Meis1* (Meis homeobox 1) is nearly ubiquitously expressed and functions as a transcription factor. A mouse *Meis1* knock-out results in embryonic lethality with prenatal alterations in the eye and vasculature and hematopoietic defects (Hisa, Spence et al. 2004). *Meis1* contains a frame-shift elongation mutation that differs between B6 (C) and NOD (CAAAAA) strains, resulting in a shorter transcript in B6 mice (ENSMUST00000118661.8). The gene is differentially expressed in the ventral tegmental area of C57BL/6J mice given chronic cocaine vs. chronic saline (Campbell, Chen et al. 2021).

*Psme4* (proteasome activator subunit 4) is nearly ubiquitously expressed and is differentially expressed in the nucleus accumbens and hippocampus of High Responder (bHR) and Low Responder (bLR) rats (Clinton, Stead et al. 2011). *PSME4* also showed differential expression in suicide mood disorder (Sequeira, Morgan et al. 2012). The SNP (rs235903318) that differs in B6 vs CC041 strains in transcript ENSMUST00000154757.2 results in a frameshift producing a truncated protein.

Finally, we identified an amino acid duplication in the WD repeat containing planar cell polarity effector gene (*Wdpcp*) The duplication is present in CC041 and not B6 and includes a residue that is conserved and has been implicated in alcohol consumption in human studies and the hypnotic effects of ethanol exposure in Drosophila.

The availability of a rapidly evolving suite of bioinformatic resources that support systems genetics analyses of the CC and other commonly used inbred mouse strains has accelerated discovery of candidate genes and genetic variants that contribute to phenotypic variation. Our high-priority candidates can be further interrogated with multi-omic approaches, cell-based assays and genetically engineered mice. Identifying and studying genes and genetic pathways that contribute to variation in behavioral responses to psychostimulants will expand our understanding novel biological mechanisms that can be targeted in drug development efforts aimed at treating and preventing substance use disorders.

---

**Suppl Figure 1.** Single scan QTL for Day 1 (saline; black line), Day 2 (saline; blue line) and Day 3 (20 mg/kg cocaine; red line) distance moved. Genome-wide significant LOD thresholds based on 1000 permutations for *p*=0.001 (red line), *p*=0.01 (green line), *p*=0.05 (blue line), and suggestive at *p*=0.1 (black line). A peak on Chr 11 was present for all three days. A QTL peak on Chr 7 was identified for Day 2 and Day 3 and a peak on Chr 14 was present for Day 1 and Day 3. QTLs on Chr 6 were specific to non-cocaine (saline) exposure days (Day 1 and 2) and a QTL on Chr 2 was unique to cocaine exposure (Day 3).

**Suppl Table 1:** Correlation of locomotor behavior in the 3-day open field test for the low cocaine response F2 population

## Notes

### Competing Interest Statement

The authors have declared no competing interest.

