## Supplemental Table 1 for "Genetic mapping in Collaborative Cross mouse strains identifies loci that affect initial sensitivity to cocaine"

**Suppl Table 1.** Correlation of locomotor behavior in the 3-day open field test for the CC041 x B6N F2 population

| Phenotypes | | r | *p* |
|---|---|---|---|
| Day 1 | Day 2 | 0.764 | $7.2 \times 10^{-87}$ |
| Day 1 | Day 3 | 0.424 | $5.1 \times 10^{-21}$ |
| Day 2 | Day 3 | 0.518 | $4.0 \times 10^{-32}$ |
| Day 1 | Day 3 - Day 2 | 0.311 | $1.7 \times 10^{-11}$ |
| Day 2 | Day 3 - Day 2 | 0.366 | $1.2 \times 10^{-15}$ |
| Day 3 | Day 3 - Day 2 | 0.986 | 0.00E+00 |
