## Supplementary figures and images for "Genetic mapping in Collaborative Cross mouse strains identifies loci that affect initial sensitivity to cocaine"

### Supplemental Figure 1

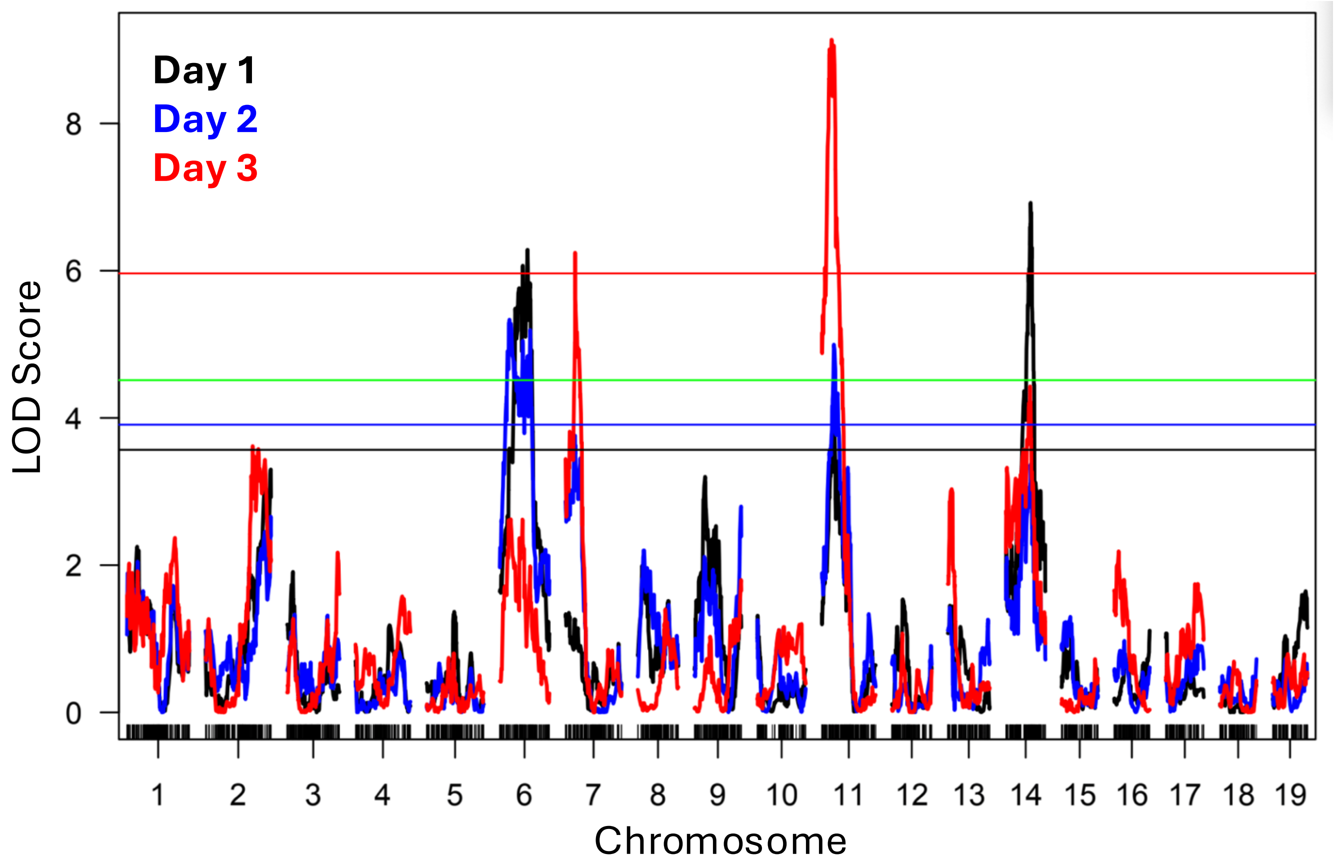
